# Management programs as a key factor for genetic conservation of small populations : The case of French local chicken breeds

**DOI:** 10.1101/2021.03.12.435064

**Authors:** Gwendal Restoux, Xavier Rognon, Agathe Vieaud, Daniel Guemene, Florence Petitjean, Romuald Rouger, Sophie Brard-Fudulea, Sophie Lubac-Paye, Geoffrey Chiron, Michèle Tixier-Boichard

**Affiliations:** Université Paris Saclay, INRAE, AgroParisTech, GABI 78350 Jouy-en-Josas, France; SYSAAF, Centre INRAE Val de Loire, UMR-BOA, 37380 Nouzilly, France; Centre INRAE Val de Loire, UMR-BOA (UR83), 37380 Nouzilly, France; Centre de Sélection de Béchanne, Hameau de Béchanne, 01370 Saint-Etienne-Du-Bois, France; ITAVI, 23 rue Jean Baldassini, 69364 Lyon Cedex 07, France

**Keywords:** Genetic diversity, local breeds, management program, conservation

## Abstract

On-going climate change will drastically modify agriculture in the future, with a need for more sustainable systems, for animal production in particular. In this context, genetic diversity is a key factor for adaptation to new conditions: local breeds are likely to harbor unique adaptive features and represent a key component of diversity to reach resilience. Nevertheless, they are often suffering from small population size putting these valuable resources at risk of extinction. In chickens, management programs have been initiated a few decades ago in France, relying on a particular niche market aiming at promoting and protecting local breeds. We conducted a unique comprehensive study of 23 French local populations, along with 4 commercial lines, to evaluate their genetic conservation status and the efficiency of management programs. Using a 57K SNP chip we demonstrated that both between and within breeds genetic diversity were high in French populations. Diversity was mainly structured according to selection and breeds’ history. Nevertheless, we observed a prominent sub-structuring of breeds according to farmer’s practices in terms of exchange, leading to more or less isolated flocks. Analysing demographic parameters as well as molecular information, we showed the efficiency of consistent management programs to conserve genetic diversity, since the earlier the breeds integrated programs the lower was the inbreeding. Finally we stressed that management programs can benefit from molecular markers and runs of homozygosity, ROH, in particular, as a valuable and affordable tool to monitor genetic diversity of local breeds which often lack pedigree information.

## Introduction

Climate change has been demonstrated to occur due to Human activities with a growing impact during the last decades (“AR4 Climate Change 2007,” n.d.). It will have multiple consequences on the availability of water and food resources but also on the distribution of diseases and of their vectors, puting many parts of the world at risk (Hoffman, 2010). This will impact the agricultural production that will have to adapt to these new harsh conditions to ensure food security, in particular in more threatened regions that often already suffer from limited food supply (Parry, 2019). Emphasis should be put on diversified and low input production models that will take advantage of locally available resources and then be less dependent on imported products (Howden et al., 2007). Indeed climate change mainly results from greenhouse gas emissions, like carbon dioxide or methane, and can be mitigated by reducing these emissions (Gerber et al., 2013). Animal breeding has been demonstrated to significantly contribute to these emissions, making domestic animals both suffering from and enhancing climate change. In addition, many ethical concerns have been raised about animal breeding, leading to the necessity to put more effort on animal welfare to fulfill consumers expectations, for example through free-range farming (Harper and Makatouni, 2002). It is thus necessary to develop a more sustainable breeding model for animals that will be able to adapt to new farming (e.g. free-range, local feed…) and climatic conditions (e.g. high temperature). In this context, genetic diversity is considered to be essential to reach this goal by conserving the adaptive potential of livestock (Notter, 1999).

Poultry production is one of the most important animal production worldwide with the highest increase during the last decade, in particular in developing countries (Windhorst, 2006). This resulted in strong selection for large-scale production systems. Indeed, following domestication, chicken populations have evolved towards a large diversity of breeds, some of them being submitted to intense selection in order to meet expectations of the market (Tixier-Boichard et al., 2011). In particular, selection of poultry led to the specialization of distinct breeds for either egg (layers) or meat (broilers) production since growth and reproduction traits are antagonistic (Fairfull and Gowe, 1990). Commercial lines, the ones under the strongest selection, exhibit a lack of rare alleles relative to ancestral populations (Muir et al., 2008). Yet, the within-breed diversity remained relatively high in broiler lines as compared to layer lines (Granevitze et al., 2007), which could be due to founder effects and different selection practices. At the same time, many traditional local breeds became at risk of disappearing, although they may still harbor original variants.

In this context, traditional local poultry breeds may represent a valuable reservoir of genetic diversity for future breeding, considering the impact that global changes could have on breeding goals regarding climate, resources or diseases (Hoffman, 2010). Yet, their population size is often limited, making them prone to suffer from inbreeding and strong genetic drift. As a result of competition against more productive breeds they are very likely to have experienced drastic and recent bottlenecks, due to a decrease of their use. Consequently, many of the local breeds are at the edge of extinction (Dávila et al., 2009). Molecular surveys have revealed that some traditional breeds retained a high diversity whereas others did not (Berthouly et al., 2008; Bortoluzzi et al., 2018; Granevitze et al., 2007). Consequently a careful management of these populations is critical to cope with the future challenges of livestock production aiming for a more sustainable breeding.

In France, local breeds have early benefited from the development of quality signs since the 50’s (Protected Origin for Bresse chicken) and the 60’s (‘Label rouge’) as documented by Verrier et al. (2005). This often consisted in high quality products with controlled process of breeding, including breed identification, free-range breeding, production in a particular area and slow growing animals with higher age at slaughter than in intensive production systems. In this context, since a few decades, initiatives have been organized to develop management programs for local poultry breeds, including pedigree recording, optimized mating plans and mild selection. These breeds are often valued locally in short chains for high quality products with a strong identification to the territory in which they are raised. They have an important patrimonial value and are often named after the name of the region or city they originated from.

In this study, we conducted a large and unique survey of 23 representative French local chicken populations to assess the impact of such management programs on genetic diversity using both molecular (57K SNP genotyping) and demographic (pedigrees and management features) data. As a comparison we also included 4 commercial lines in order to provide a comprehensive view of the diversity patterns associated with different kinds of management. This assessment makes it possible to draw some lessons and make recommendations for the management of genetic diversity that could be extended to any conservation program.

## Material and methods

### Material

The 23 local breeds were chosen in order to well represent the diversity of origins and localities of French breeds. The sampling strategy involved three situations of breed management. Group 1 involved 19 French local breeds which were engaged in a management program aimed at marketing a high quality product while preserving the genetic diversity and the originality of each breed. Sampling of animals took place in a pedigreed nucleus flock set up in the Breeding Center of Béchanne in 2013. Group 2 involved 3 local breeds not yet engaged in such a program, so these animals were sampled from 2 to 6 independent farmers, depending on the breed, from 2013 to 2014. In addition, one breed, the Marans, was represented in both groups, with 3 lines maintained by a single breeder that were included in Group 1 (breed code MAG), and different color varieties maintained by six fancy breeders motivated by the conservation of the breed, included in Group 2 (breed code MAR). Local breeds information is summarized in table 1. Finally, Group 3 included 4 commercial lines as control populations to assess breed identity and possible introgression events: DNA samples of two fast growing broiler lines (FG1 with 42 individuals and 43 for FG2) were obtained from the DNA collection set up for the Aviandiv EU project in 1999 (Hillel et al., 2003) to be genotyped in this study; genotyping data were obtained for one line of French slow growing high quality ‘label’ chicken, SGL (96 individuals) and one line of brown-egg layer, LAY (57 individuals). A data transfer agreement was established with each owner of these lines in order to re-use existing genotyping data without re-sampling but phenotypic data were not provided.

**Table 1.** Number of animals genotyped (Nb_anim), mean geographic location, history of creation and main morphological features for French local breeds, either engaged in a management program (Group 1) or distributed among fancy breeders (Group 2)

| Breed name | Breed code | Group | Nb_anim | Geographic area in France | Center of origin mean coordinates | Historical records | Plumage color |
| --- | --- | --- | --- | --- | --- | --- | --- |
| Poule d'Alsace | ALS | 1 | 30 | North East | 48.57 N ; 7.75 E | 19th century | black |
| Barbezieux | BAR | 1 | 59 | South-West | 45.47 N ; -0.15 W | 19th century | black |
| Bourbonnaise | BOU | 1 | 57 | Centre | 46.57 N ; 3.34 E | n.a. | silver |
| Bresse Gauloise Blanche 11 | B11 | 1 | 60 | Centre-East | 46.21 N ; 5.23 E | ancestral Bresse (16th century) | white |
| Bresse Gauloise Blanche 22 | B22 | 1 | 60 | Centre-East | 46.21 N ; 5.23 E | Created in 2000 from B11 | white |
| Bresse Gauloise Blanche 55 | B55 | 1 | 43 | Centre-East | 46.21 N ; 5.23 E | 1950 (egg production) | white |
| Charollaise | CHA | 1 | 56 | Centre-East | 46.44 N ; 4.28 E | 1950 (meat) | white |
| Contres | CON | 2 | 41 | Centre | 47.42 N ; 1.43 E | 20th century | silver |
| Cou nu du Forez | CNF | 1 | 59 | Centre-East | 45.74 N ; 4.23 E | n.a. | white |
| Coucou de Rennes | CDR | 1 | 55 | West | 48.11 N ; -1.68 W | 19th century | Sex-linked barring |
| Gasconne | GAS | 1 | 60 | South-West | 43.43 N ; 0.58 E | 19th century | black |
| Gâtinaise | GAT | 1 | 57 | South of Paris | 48.15 N ; 2.70 E | 5th century | white |
| Gauloise Grise | GG | 1 | 60 | Centre-East | 46.21 N ; 5.23 E | 16th century | Autosomal barring |
| Gauloise noire | GN | 1 | 58 | Centre-East | 46.63 N ; 5.22 E | 16th century, currently bred for egg production | black |
| Geline de Touraine | GDT | 1 | 59 | Centre-West | 47.13 N ; 1.00 E | 15th century | black |
| Gournay | GOU | 1 | 58 | West | 49.48 N ; 1.72 E | 19th century | mottled |
| Grise du Vercors | GDV | 1 | 44 | South-East | 45.02 N ; 5.29 E | Created in 2002 from a cross | Sex-linked barring |
| Houdan | HOU | 1 | 58 | West of Paris | 48.79 N ; 1.60 E | 17th century | mottled |
| Hergnies | HER | 2 | 60 | North | 50.48 N ; 3.52 E | Recent reconstitution | Autosomal barring |
| Le Mans | MAN | 2 | 29 | Centre-West | 48.01 N ; 0.20 E | Recent reconstitution | black |
| Marans managed | MAG | 1 | 52 | Centre-West | 47.62 N ; -1.19 W | Commercial breeding for eggs | Brown red, red, silver cuckoo, |
| Marans fancy | MAR | 2 | 65 | Centre-West | 46.3 N ; -1.0 W | Fancy breeders, first described in 15th century | Black, Brown red, red, silver cuckoo, white, wheaten, wild-type, |
| Merlerault | MER | 1 | 38 | West | 48.7 N ; 0.29 E | 19th century | black |
| Noire du Berry | NDB | 1 | 56 | Centre | 46.95 N ; 1.99 E | 19th century | black |

The number of individuals studied per population was 56 on average, and ranged from 29 for the MAN breed to 96 for the ‘label’ chicken line SGL, for a total of 1512 individuals. A blood and DNA collection for 1350 animals of the 24 local populations (Group 1 + Group 2 and the two Marans breeds MAG and MAR) was stored under the project name BioDivA at the @BRIDGe biological resource center of CRB-Anim infrastructure (CRB-Anim, 2018), and a material transfer agreement was signed with each breed’s association or each individual farmer. Family structure was avoided, by sampling distinct families as much as possible.

Local breeds were described by their geographic origin, morphological standard, historical records (Table 1). Parameters of the management program include number of founders, either males or females, number of generations since the start of the program with pedigrees (Table 2).

**Table 2.** Pedigree data available on breeds from group 1 except Marans* (MAG), genetic size (Ne_demo) calculated according to Cervantes et al (2011), mean inbreeding coefficient at sampling, mean number of chicks produced each year.

| Breed | Number of male founders | Number of female founders | First year with pedigree | Number of generations in 2013 | Number of sires in 2013 | Number of dams in 2013 | Mean inbreeding coefficient at sampling (%) | Pedigree based Ne in 2013 (Ne_demo) | Initial Founders based Ne (Ne_found) | Inbreeding rate per generation in % (Delta_F) |
| --- | --- | --- | --- | --- | --- | --- | --- | --- | --- | --- |
| Poule d'Alsace (ALS) | 20 | 33 | 2008 | 5 | 27 | 74 | 2.3 | 65 | 49.8 | 0.46 |
| Barbezieux (BAR) | 7 | 7 | 2002 | 11 | 52 | 116 | 7.0 | 27 | 14 | 0.64 |
| Bourbonnaise (BOU) | 75 | 232 | 2005 | 8 | 40 | 100 | 2.4 | 114 | 226.7 | 0.30 |
| Bresse Gauloise Blanche B22 | 26 | 102 | 2000 | 13 | 99 | 215 | 5.3 | 108 | 82.9 | 0.41 |
| Bresse Gauloise Blanche B11 | 23 | 256 | 1989 | 24 | 105 | 346 | 7.1 | 122 | 84.4 | 0.30 |
| Bresse Gauloise Blanche B55 | 23 | 255 | 1989 | 24 | 104 | 376 | nd | 153 | 84.4 | NA |
| Charollaise (CHA) | 34 | 103 | 2005 | 8 | 20 | 43 | 2.7 | 63 | 102.3 | 0.34 |
| Cou nu du Forez (CNF) | 34 | 98 | 2007 | 6 | 20 | 50 | 3.5 | 53 | 101 | 0.58 |
| Gasconne (GAS) | 5 | 5 | 2009 | 4 | 24 | 77 | 7.2 | 22 | 10 | 1.80 |
| Gâtinaise (GAT) | 11 | 12 | 2006 | 7 | 40 | 112 | 5.6 | 56 | 23 | 0.80 |
| Gauloise Grise (GG) | 77 | 252 | 1998 | 15 | 65 | 184 | NA | 285 | 235.9 | NA |
| Gauloise Noire (GN) | 108 | 275 | 1996 | 17 | 20 | 83 | 5.3 | 119 | 310.2 | 0.31 |
| Géline de Touraine (GDT) | 43 | 103 | 1996 | 17 | 55 | 110 | 4.2 | 142 | 121.3 | 0.25 |
| Gournay (GOU) | 24 | 61 | 2003 | 10 | 46 | 104 | nd | 98 | 68.9 | 0.45 |
| Grise du Vercors (GDV) | 29 | 83 | 2007 | 16 | 24 | 53 | 2.7 | 65 | 86 | 0.17 |
| Houdan (HOU) | 30 | 81 | 2004 | 9 | 24 | 58 | NA | 80 | 87.6 | NA |
| Merlerault (MER) | 18 | 32 | 2006 | 7 | 14 | 34 | 5.6 | 44 | 46.1 | 0.8 |
| Noire du Berry (NDB) | 27 | 47 | 2008 | 5 | 47 | 103 | 2.7 | 99 | 68.6 | 0.54 |

NA : not available
\* : pedigree data were not available for the Marans breed from group 1 (code MAG) which was managed by a commercial breeder

### Methods

Using management program information we computed effective population sizes either estimated i) at the start of the program from the number of founders and their sex-ratio (Ne_found) using the classical formula, 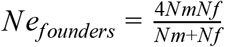, with Nm and Nf being the number of sires and dams respectively, or, ii) at sampling date (2013) using complete pedigrees (Ne_demo) according to (Cervantes et al., 2011) for breeds maintained by the Bechanne breeding center (Group 1). Individual inbreeding coefficients, F_pedig, were also estimated using the pedigree. Considering that coancestry and inbreeding were null at the start of the pedigree, dividing F_pedig by the number of generations in the pedigree we estimated mean inbreeding rate, Delta_F. Pedigree data were collected from the SYSAAF database with the agreement of each breed’s association.

Genotypes were obtained using the 57K Illumina Beadchip (Groenen et al., 2011). After mapping on the current version of chicken genome assembly (Gal_gal_6), we kept 51041 SNPs well located and identified. We then applied quality control filters using Plink 1.9 software (Chang et al., 2015) with a minor allele frequency (MAF) larger than 0.001 (*--maf* function), and call rates of at least 5% for both markers and individuals (*--mind* and *--geno* functions). This resulted in a set of 46940 autosomal SNP for 1512 individuals with a total genotyping rate of 99.86%.

For some analyses we also constructed a pruned dataset for which we removed linked markers with Plink 1.9 software by applying the *--indep-pairwise* function with a window of 30 SNP, sliding by step of 10 SNPs and a r^2^ threshold of 0.8 above which markers are considered linked. It resulted in a 45209 autosomal SNPs dataset.

The level of diversity at the population level was investigated through 5 different summary statistics all computed with Plink 1.9 software: i) The mean individual inbreeding coefficient computed using allelic frequencies estimated over all populations, ii) the mean individual inbreeding coefficient computed using allelic frequencies estimated for each population, iii) the mean observed heterozygosity, iv) average MAF, and v) the proportion of fixed alleles in the population. The two first indices were computed using the method of the moment (Purcell et al., 2007) on the pruned dataset (*--het* function) and were very similar to Fit and Fis respectively (under the assumption that deviation to Hardy-Weinberg was only due to inbreeding) and thus will be named like this hereafter. Global Fst’s were also computed on pruned dataset between populations and groups of populations according to breeders or lines (*--fst* function).

Runs of homozygosity (ROH) were computed using the *--homozyg* function of Plink 1.9 software with the following parameters: minimum size of 500kbp and minimum number of 30 SNP, a density of at least 1 SNP every 50kbp, allowing for 1 missing and heterozygous SNP per window of a length of 50 SNP). The proportion of ROH in the genome was turned into inbreeding coefficients according to McQuillan et al., (2008).

Genetic effective population sizes were computed according to the level of linkage disequilibrium (Ne_LD hereafter) using Ne-Estimator v2.1 software (Do et al., 2014). Since this approach is very sensitive to physical linkage between markers we applied a drastic thinning to the autosome dataset in order to keep one SNP over 500. This led to 94 autosomal SNPs either in different chromosomes or with a minimal distance of 3.9 Mbp from each other when located on the same chromosome which is sufficient to ensure for no physical linkage even in pure chicken lines (Fu et al., 2015).

Genetic distances between all individuals were computed as an identity by state distance matrix (*--distance square ibs* function of plink 1.9). This matrix was then used to compute and plot an unrooted neighbor-joining tree using the APE R package (Paradis and Schliep, 2019). The trees plotting for MAR and HER populations were made using the ggtree R package (Yu et al., 2017) in order to figure out additional information such as the breeders and the phenotype.

Pairwise genetic distances between populations were computed as Nei’s distance D or Fst using the Stampp R package (Pembleton et al., 2013). The Nei’s pairwise D distances were then used to compute a neighbor-net tree of populations using Splitstree software (Huson and Bryant, 2006). Correlation between pairwise Fsts and geographical distances were tested with a Mantel test using the Vegan R package (Dixon, 2003).

In a previous study of Berthouly et al (2008), 14 French local breeds were separated into two groups according to their genetic similarity with asiatic breeds, as a result of importation of Asiatic breeds in the 19^th^ century. In order to search for this structure in the current study, we investigated population structure using a Discriminant Analysis on Principal Components, DAPC, using the ADEgenet R package (Jombart, 2008; Jombart et al., 2010) with a number of clusters K set to 2. The probability for a given breed to belong to each cluster was plotted according to their geographic origin (French departments) in order to reveal possible centers of dissemination following introduction of asiatic breeds and plotted on a map along with sampling locations.

All additional computations and graphical representations were done using R (R Core Team, 2019) and ggplot2 (Wickham, 2011) and corrplot (Wei et al., 2017) R packages.

## Results

### Demographic parameters

The Ne_demo values ranged from 22 to 285 in 18 breeds from Group 1 (Table 2). Interestingly, the populations with the lowest values of Ne_demo exhibited a relatively recent management program. The mean year of the onset of the program was 1996 for breeds with Ne_demo above 100 whereas it was 2006 for breeds with Ne_demo below 100. Considering that the generation interval is generally 1 year, the populations with the highest number of generations tended to exhibit a large Ne_demo, most probably because their number of founders was generally higher than for recently managed breeds. The mean Ne_found at the start of the program was 100 but varied greatly between populations from 10 for the GAS breed to 310 for the GN breed (table 2). Ne_demo increased as compared to Ne_found for 11 populations, particularly in Gauloise Grise and Bresse B55, but decreased for 7 populations, particularly in Gauloise noire and Bourbonnaise. The mean pedigree based inbreeding coefficient at sampling was mild, with 4.5% and exhibited a narrow range of variation (from 2.3 to 7.2). The ratio of female to male founders generally varied from 1 to 4, except for two populations (Bresse Gauloise blanche B11 and B55) where it reached 11.

### Molecular information

#### Genetic diversity

The genetic diversity within populations was characterized by the mean of 5 indices (figure 1; Sup. Mat. S4). Mean Fit ranged from 0.11 for the NDB breed to 0.46 for the B22 breed with a mean value of 0.26 (standard error 0.09). Mean Fis ranged from −0.03 for the BOU breed to 0.08 for the HER breed with a mean value of −0.002 (s.e. 0.03). The mean observed heterozygosity rate was 0.34 (s.e. 0.01) and ranged from 0.31 for the MAG breed to 0.36 for the B55 breed. The mean MAF across breeds was 0.21 (s.e. 0.03) and ranged from 0.15 for B22 breed to 0.25 for NDB breed. Finally the mean proportion of fixed alleles was 0.18 (s.e. 0.09) and ranged from 0.05 for the MAR breed to 0.38 for the B22 breed. The mean Ne_LD across populations was about 239 (sd 457.5) with a minimum value of 2.5 for the MAN breed and a maximum value of 1840 for the B11 population (Sup. mat. S4).

**Figure 1.**
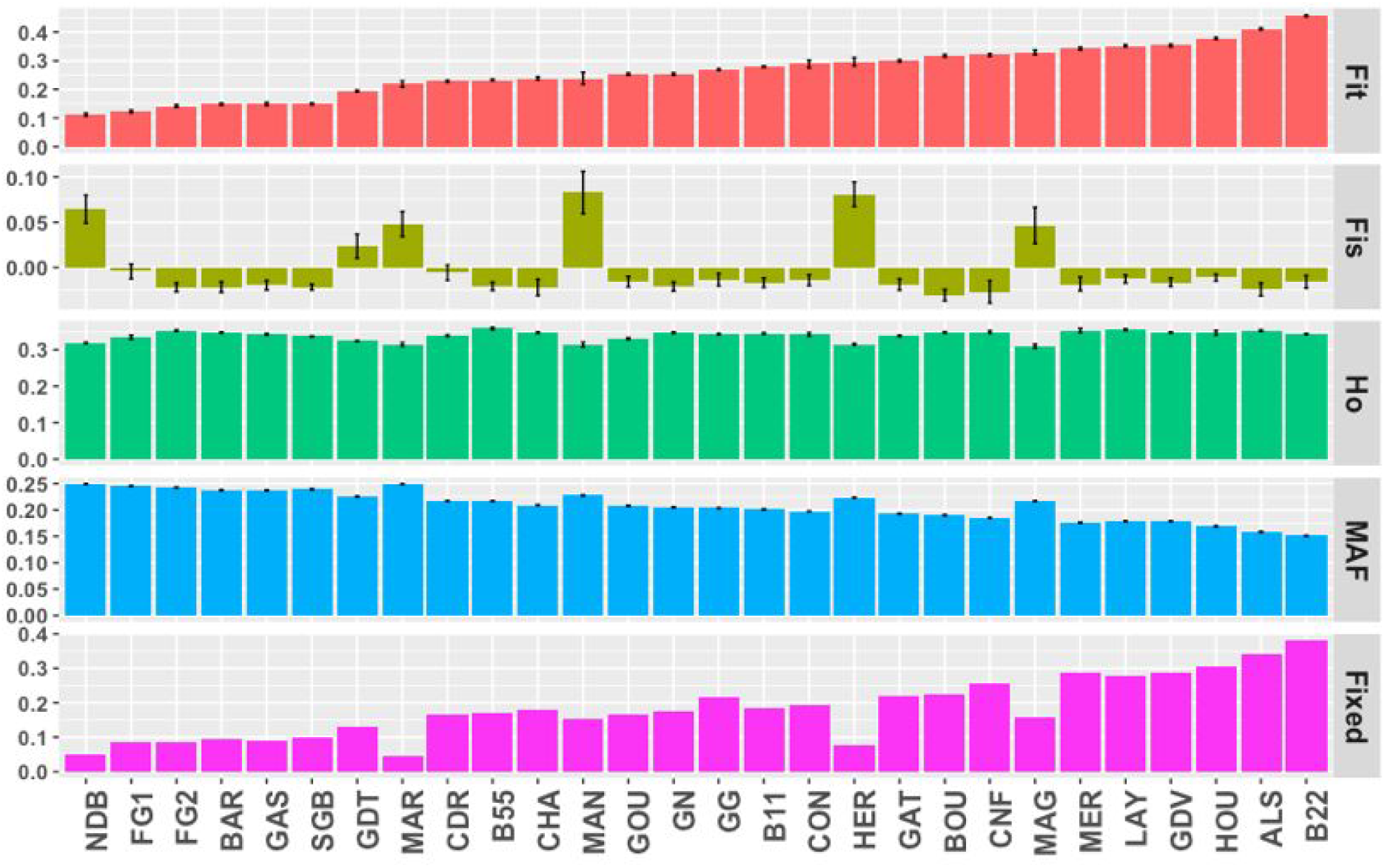
Genetic diversity summary statistics for each breed, with from the top to the bottom Fit, Fis, Observed heterozygosity (Ho), Minor Allele Frequency (MAF) and the proportion of fixed alleles (Fixed). Populations were sorted according to the Fit.

#### Runs of homozygosity

Runs of homozygosity (ROH) have been detected and turned into inbreeding coefficients, F-ROH (figure 2; Sup. Mat. S4). These mean F-ROH per population ranged from 0.13 for NDB to 0.42 for B22 with a mean over all breeds of 0.24 (s.e. 0.01). The mean length of homozygous segments per population was 2.9 Mb (s.e. 112 kb) and it ranged from 2.2 Mb for the B55 breed to 4.7 Mb for the MAN breed. The mean number of ROH per individual per population was 76.30 (s.e. 4.51). It ranged from 36.58 for the NDB breed to 128.40 for the B22 breed.

**Figure 2:**
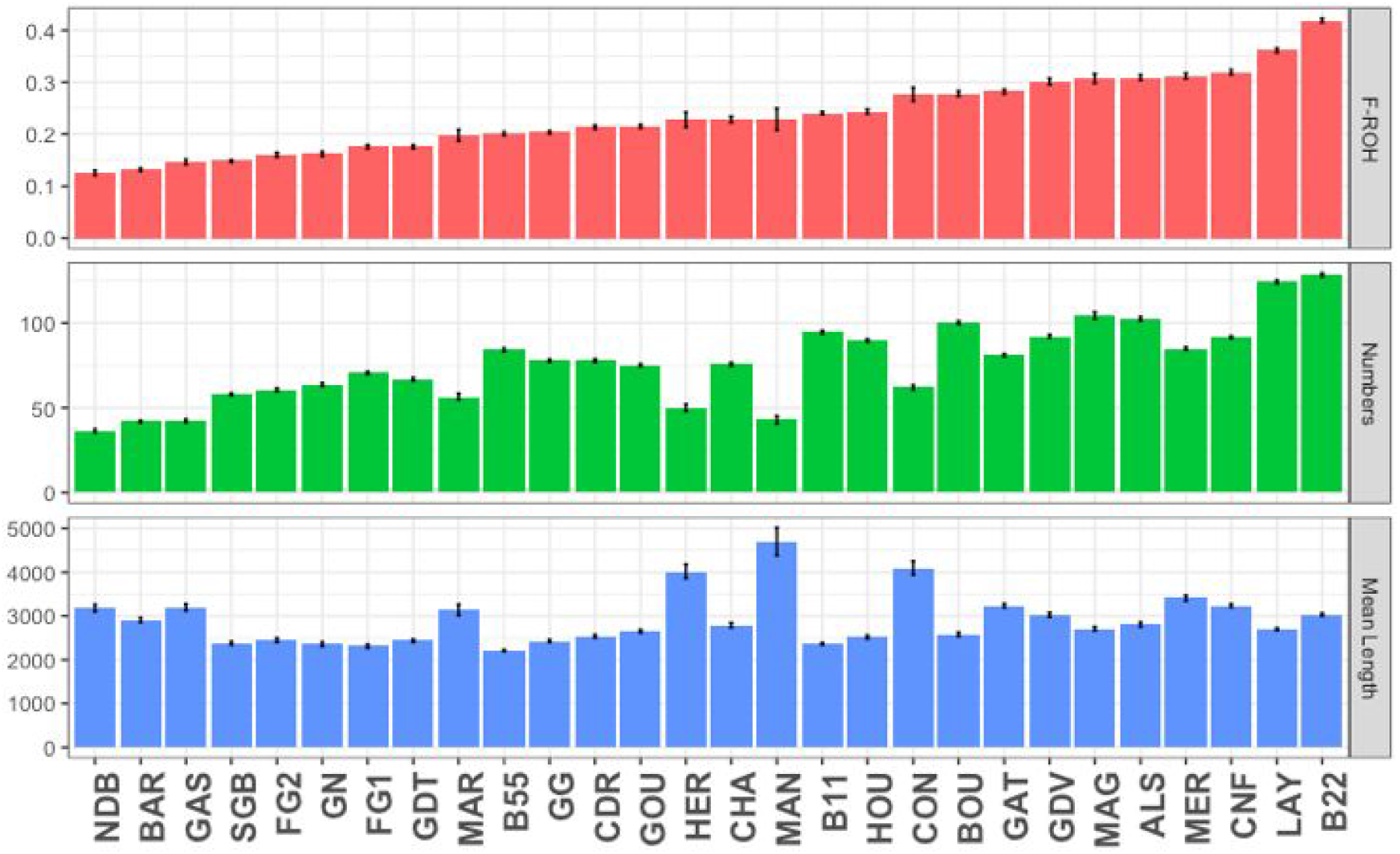
Runs of Homozygosity (ROH) summary statistics per population with from the top to the bottom, the inbreeding coefficient (F-ROH), the mean number of ROH per individual and the mean length of each ROH. Populations were sorted according to the F-ROH.

All correlations between molecular and demographic estimates of genetic diversity for the group 1 are all presented in the figure 6.

#### Genetic structure

### All populations

All populations were almost well separated genetically and clearly distinguishable from each other (Figure 3). The MAG population had three distinct groups with the LAY population inserted in-between. Similarly the HER population was separated into several groups among which the GG population was inserted.

**Figure 3.**
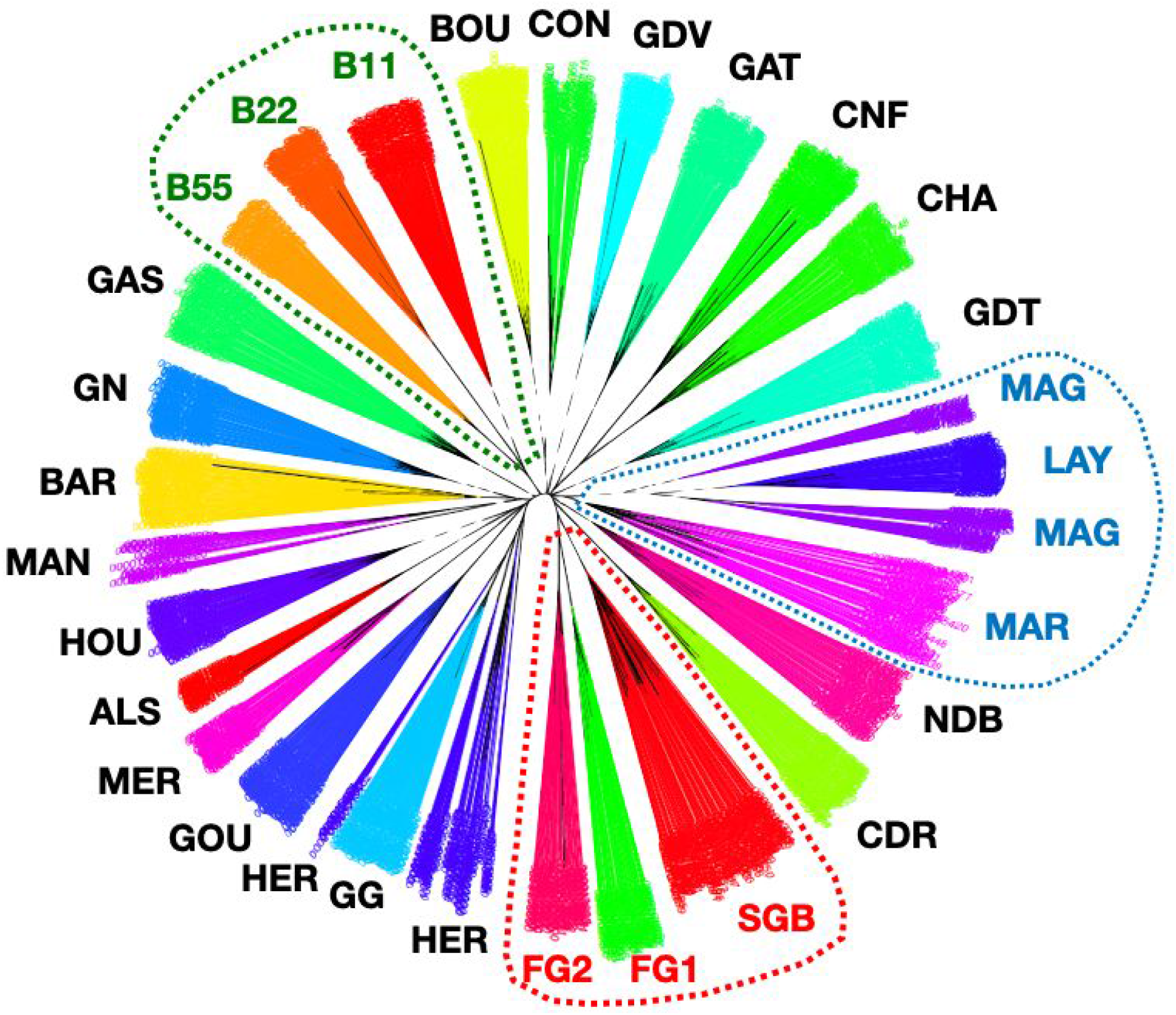
Unrooted neighbor joining tree of individuals with respect to populations. Dot lines stand for grouping features: Brown egg laying breeds (blue), broiler breeds (red) and Bresse breed (green).

**Figure 4.**
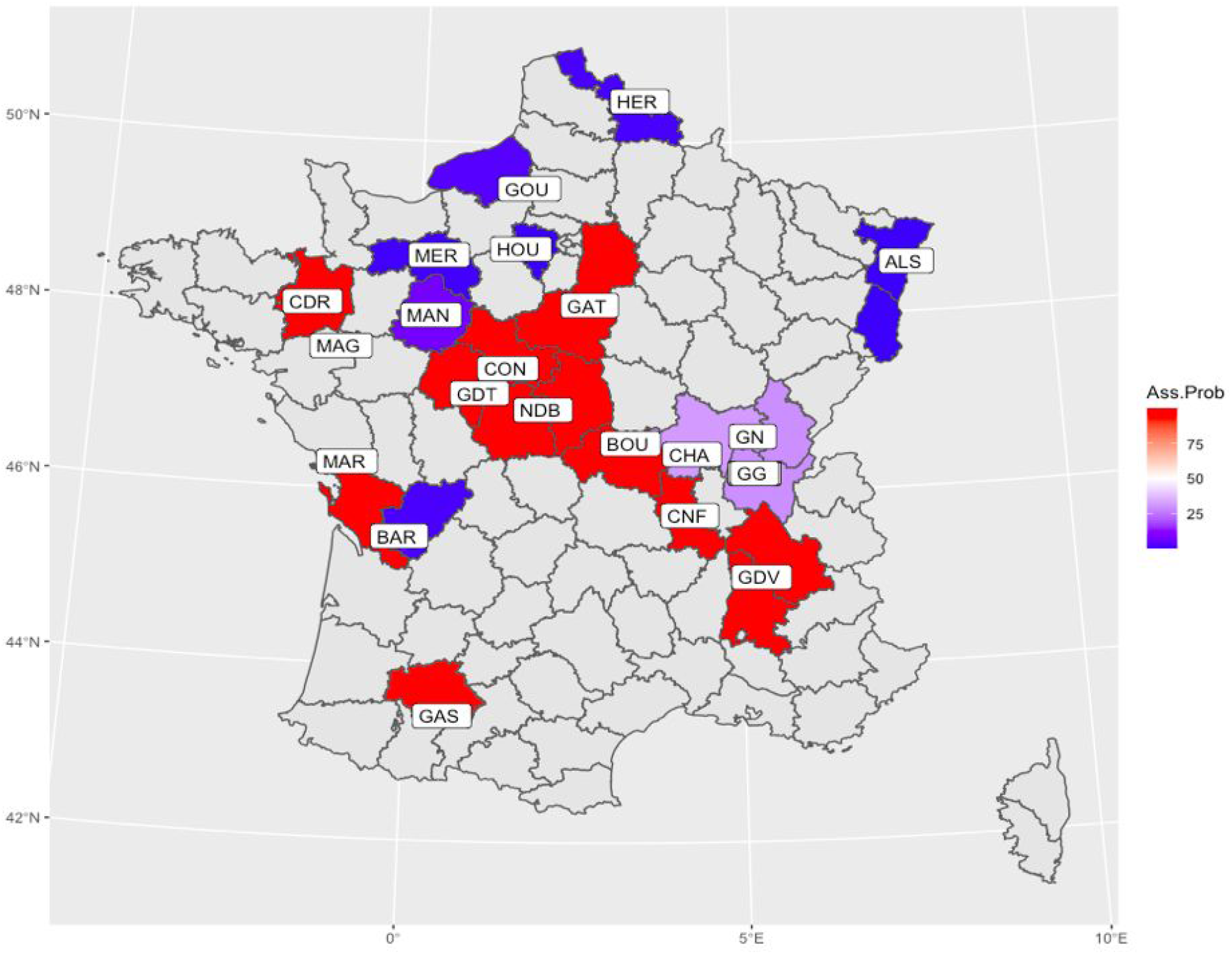
Mean center of origin of breeds and assignation of French regions (departments) according to breed geographical distribution and assignment of the populations to each of the two clusters of the DAPC, either Asiatic (red) or European (blue).

The mean weighted Fst between all populations was 0.26. Pairwise Fst were also estimated and were all significantly different from 0 (Supplementary material, S1). They ranged from 0.12 between MAR and NDB populations to 0.44 between LAY and B22 populations.

The mean assignment probability to each of the 2 clusters in the DAPC was averaged for each population (Supplementary material S3) which showed a clear affiliation of each population to a given cluster except for the B55 population. Figure 5 showed the geographic location of breeds together with the most probable cluster they belong to. The neighbornet tree (Figure 5) confirmed the separation between the two clusters of origin, European or Asian, except in the case of the 3 Bresse populations (B11, B22, B55) which grouped together on the edge of the Asian cluster, although B11 and B22 lines showed a higher probability of assignment to the European cluster. In addition, we observed a close relationship between the MAG and LAY populations on the one hand and between the HER and GG populations on the other hand.

**Figure 5.**
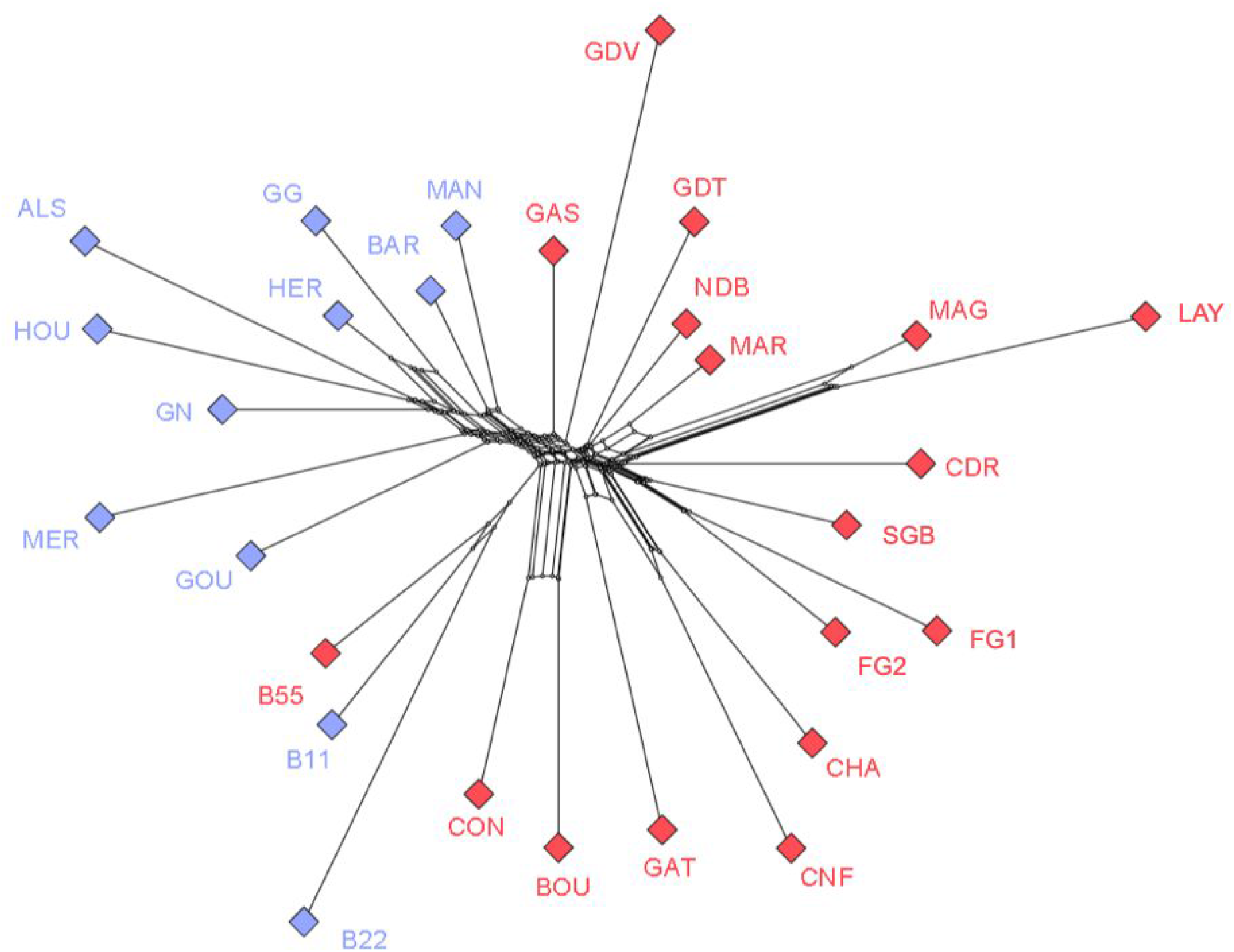
Neighbornet tree of the 28 chicken populations with respect to their cluster affiliation, red for Asian and blue for European.

**Figure 6.**
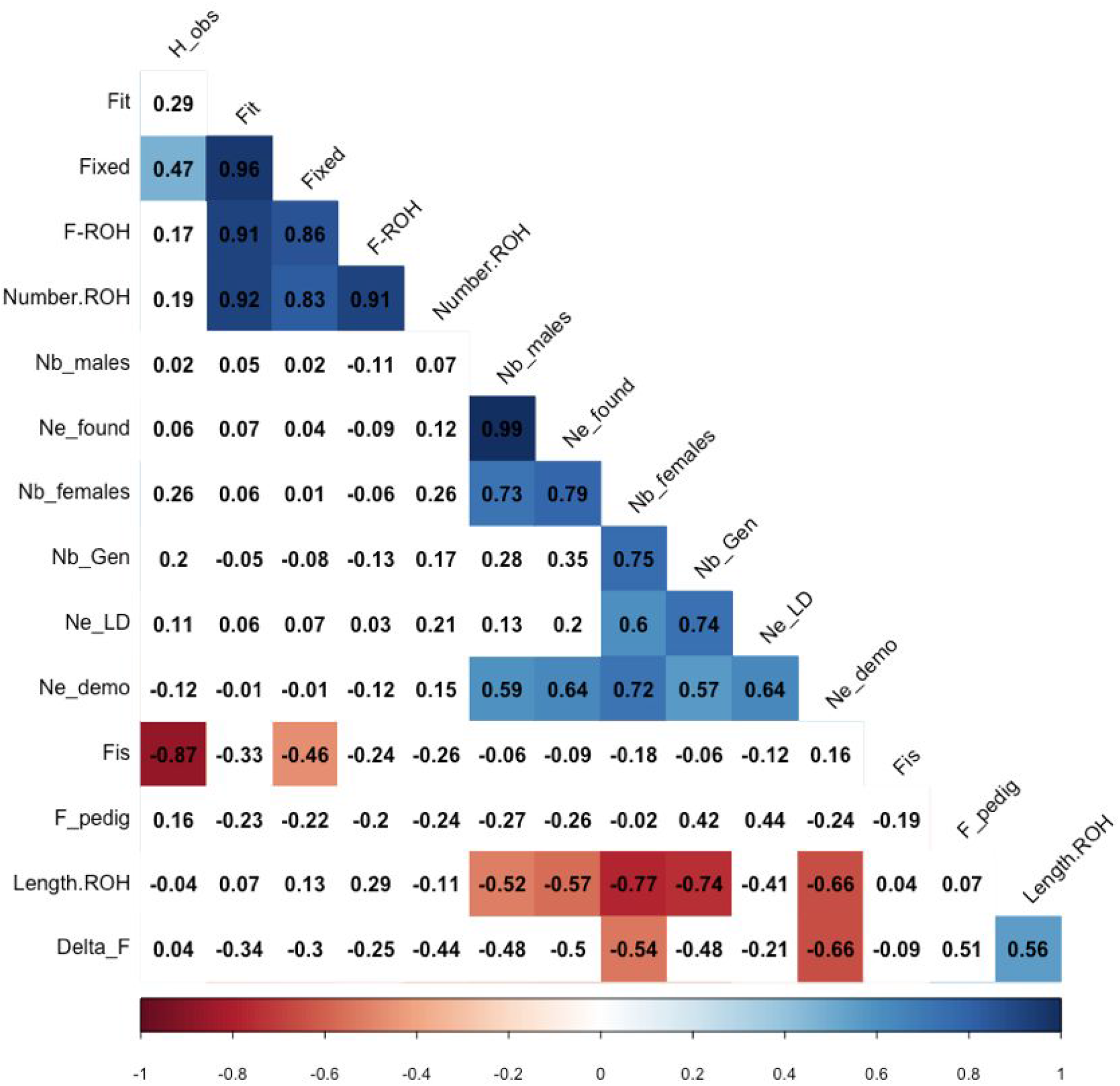
Correlation matrix of molecular or demographic diversity indices for breeds undergoing a management program (Group 1). Values correspond to Pearson’s correlation coefficients. Colored cells stand for significant correlation (p<0.05) either positive (blue) or negative (red), intensity depending on the strength of the correlation between estimates. White cells represent non-significant correlations.

The broiler populations were grouped together, fast growing populations (FG1 and FG2) being closer to each other than to the slow growing ‘label’ population (SGL).

Mantel test of the correlation between *1/(1-Fst)* and the log of geographical distance was not significant.

### Within breed genetic structure

Four breeds exhibited a within population structure, the Marans (MAR & MAG), HER, MAN and CON. This genetic structure is mainly due to a breeder effect that will be detailed further for Marans and HER for which we had a higher number of breeders (table 1).

The MAR (Group 2) and the MAG (Group 1) subpopulations of the Marans breed were clearly separated whereas the LAY population (Group 3) was surprisingly intermingled in the MAG group, where different sublines could be distinguished. The Fst between MAG and MAR clusters was 0.16. The global Fst between all 7 breeders was 0.15 while the Fst between the 6 breeders of the MAR group was 0.06. Another clustering was also visible for the Marans breed according to the phenotype, namely feather color, for which we observed a Fst of 0.17. While considering MAR and MAG apart, we observed a between-colors Fst of 0.12 (6 colors) and 0.20 (3 colors) respectively. This genetic structure was presented in figure 7.

**Figure 7.**
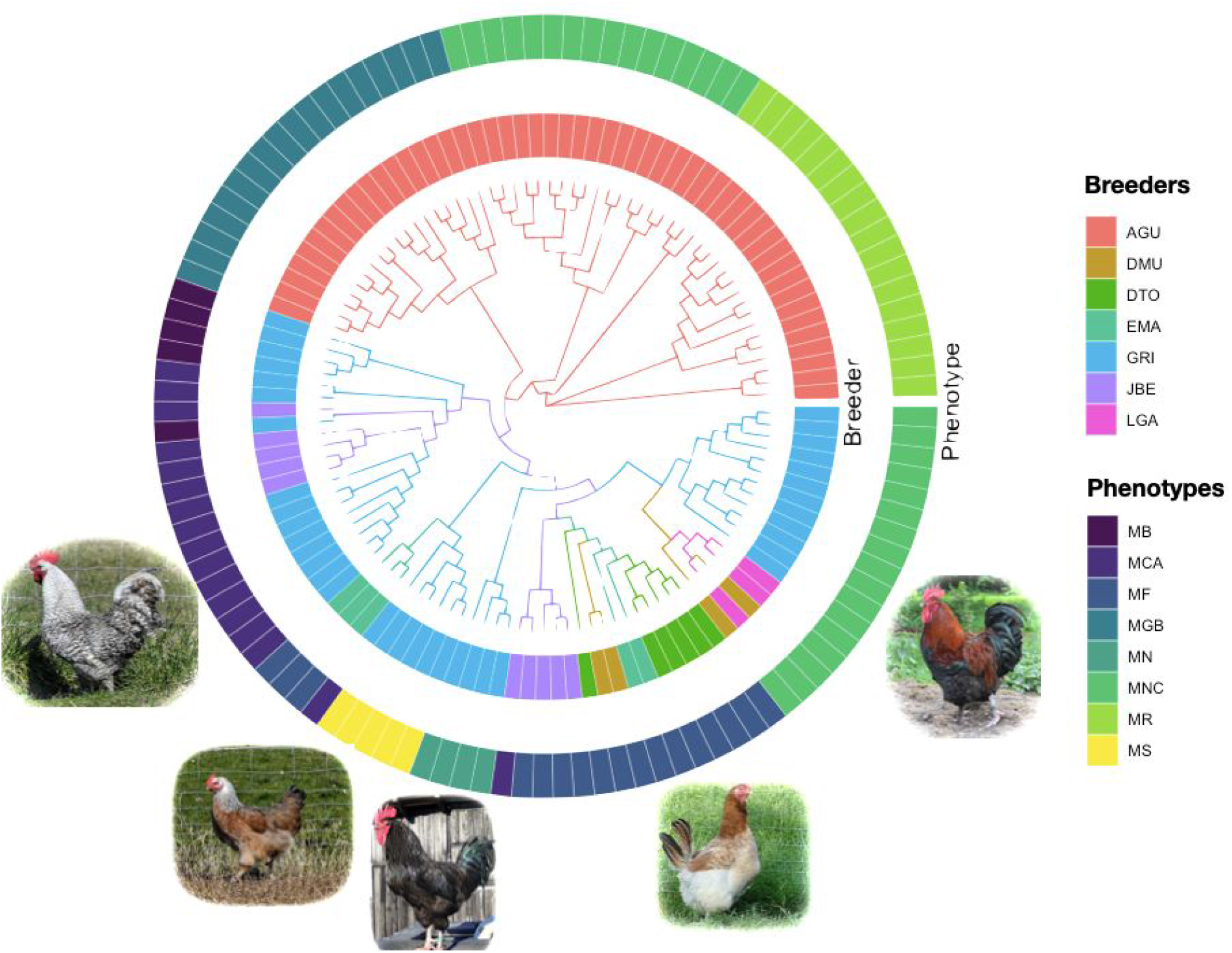
Unrooted neighbor-joining tree of the Marans breed. Colors of the surrounding circles represent the breeders (inner circle) and the phenotypes (outer circle).

The Hergnie breed, HER, was also strongly structured according to the 6 fancy breeders with a corresponding Fst of 0.20 (Supplementary material, S2). The CON and MAN breeds also exhibited a genetic structure according to the fancy breeders with a Fst of 0.11 between the 3 CON flocks and of 0.22 between the 2 MAN flocks.

## Discussion

Molecular markers have boosted the analysis of genetic diversity and its spatial or temporal dynamics. Yet, parameters in the hands of population managers are most often demographic, such as number of families, mating plan, exchange of breeding animals. Whereas both types of data are generally available for selected populations of livestock, this is rarely the case for local breeds. We will first discuss the information brought by molecular data on the genetic diversity of local chicken breeds and address the special situation of the Marans breed, which illustrates different types of management and includes feather color, a phenotype generally considered in priority by amateur breeders. Finally, we will discuss the usefulness of molecular tools and their relationship with the demographic parameters in order to make recommendations for launching conservation programs of local breeds.

### Genetic structure of French local breeds

#### Overall breed structure

French chicken breeds exhibit a large genetic diversity with breeds clearly differentiated from each other. The high level of genetic differentiation between breeds with an overall Fst of 0.26 was comparable to what was observed in other European countries, with values of 0.25 for British breeds (Wilkinson et al., 2012) or 0.22 for Hungarian (Bodzsar et al., 2009). It is even higher than a previous study conducted on 14 French breeds in which they observed an overall Fst of 0.19 (Berthouly et al., 2008). However the observed genetic differentiation between French breeds is larger than the one observed in Chinese populations with overall Fst of 0.11 (Qu et al., 2006) and pairwise Fst ranging from 0.03 to 0.27 (Zhang et al., 2020) while we observed values from 0.13 to 0.43 for French local breeds. This indicates that management of French local breeds succeeded in conserving a large number of distinct breeds.

#### Effect of selection and usage

Despite clear separations between breeds, we observed proximities that can be explained by selection, history or life history traits. Indeed the breeds mainly dedicated to egg production, i.e. brown egg layers, were grouped together whatever their status, either commercial (LAY) or local, managed or not (MAG and MAR respectively). Similarly the meat producing breeds, i.e. broilers, clustered together whatever being slow (SGL) or fast growing breeds (FG1 and FG2). However the different lines of the Bresse breed (B11, B22 and B55), an emblematic French free-range slow growing breed, clustered together but were apart from the 3 commercial broilers control breeds. This is probably due to the fact that a large majority of French local breeds were initially dual-purpose breeds even if they are now mainly raised for their meat and are poor egg producers. Selection pressure on growth rate was weak and mainly consisted in stabilizing growth rate and adult body weight, leading to a lack of clustering of the Bresse breeds with the other broiler breeds. Indeed management of French local breeds is done with moderate emphasis on quantitative production traits while maintaining the breed standard in terms of body conformation and phenotypes (e.g. plumage and comb).

#### Spatial structure

Geographical proximities between center of origin of breeds also appeared to structure the genetic diversity of the breeds at a fine scale. Indeed the CNF and CHA breeds, both originating from the same French region were genetically close together, as well as the MER, HOU and the GOU breeds. This illustrates the historical exchanges between close farmers in markets but also the creation of the breeds often based on the crossing of multiple populations locally available as for CNF which originated from both CHA and GAT. Nevertheless we did not find any signal of isolation by distance at the overall level. This is probably due to multiple introductions and origins (Denis, 2017). In addition the clear structuring of the French breeds into two groups is in accordance with Asian and European origins previously described (Berthouly et al., 2008). This is consistent with the history of chickens in Europe, characterized by an initial dispersion from Asia early on, around the Iron Age, followed by a second introduction of Asian breeds 150 to 200 years ago (Malomane et al., 2019). The presence of these two groups can be of interest for selection perspectives if heterosis can be observed between these two groups, making them heterotic groups which are widely used in plant breeding (Brandenburg et al., 2017). Further insights with crossing experiments are needed to confirm the interest of this possible future use of local chicken breeds.

### Within breed genetic diversity

#### Overall diversity

The genetic diversity of the French breeds was globally high with a mean within breed observed heterozygosity of 0.34 and ranged from 0.31 to 0.36. These values are larger than those observed in other European breeds with an observed heterozygosity 0.19 in breeds sampled in Germany (Malomane et al., 2019) and observed heterozygosity ranging from 0.16 to 0.29 in Dutch breeds (Bortoluzzi et al., 2018). Reversely this level of within population diversity is very close to the one observed in Asian populations with values ranging from 0.29 to 0.36 (Zhang et al., 2020). Interestingly this corresponds to the level of diversity observed in populations less structured like the Chinese ones in which gene flow occurs regularly between breeds ensuring the conservation of a high level of within-breed genetic diversity. On the contrary, other European populations exhibit strong genetic structure with limited gene flow between breeds but limited within breed diversity due to very small populations size, often maintained by fancy breeders without a proper conservation program. Here, the French populations exhibit a high level for both within and between breeds genetic diversity. This is the result of initiatives to conserve numerous breeds favored by policies supporting niche markets of ‘label’ products coupled with effective management of breeds through conservation or low intensity selection programs. Indeed, most Fis values were close to 0 (−0.002), which is congruent with a previous study (Berthouly et al., 2008) indicating no deficit of heterozygosity. Reversely, in other European countries local breeds exhibited higher Fis values, as observed in Great Britain (Wilkinson et al., 2012), Germany (Malomane et al., 2019) or The Netherlands (Bortoluzzi et al., 2018), where these breeds are mainly raised for ornamental purposes. Our results thus suggest a good monitoring of inbreeding through precise mating rules based on pedigree records in order to limit coancestry. This was confirmed by the breeds from group 2, in particular HER, CON and MAN, that are not involved in a management / conservation program and exhibited longer ROH than breeds from group 1 involved in management programs. Even if inbreeding estimates based on ROH were relatively moderate in group 2, this indicated recent inbreeding probably due to fragmentation into disconnected flocks.

#### Sub-structuring of breeds

Some breeds, Marans (including both MAR and MAG), MAN, HER and NDB, exhibited high values of Fis. All of them except MAG, do not benefit from a conservation program but are raised in multiple flocks due to different breeders or varieties within breeds (ie group 2). Thus, these large Fis values likely result from a Wahlund effect due to sub-structuring of breeds. This was confirmed with a weak level of fixed alleles and larger minor allele frequencies at the population level, due to conservation of numerous alleles in different flocks with limited gene flow between them. For the MAG and the HER breeds we observed close proximity with LAY and GG respectively. For MAG breed this could suggest the use of commercial layer individuals at the start of the breeding program to improve performance. After investigation on the field we discovered that the French local GG breed was used by one amateur HER breeder to improve the phenotype and the genetic diversity of his flock. Indeed this latter breed is only raised by a very few number of breeders and is thus threatened. Reversely the MAG breed was raised very professionally with more emphasis toward production in a particular market.

#### The case of the Marans breed

The Marans breed, including MAR and MAG, is a very interesting case study since it encompasses numerous varieties mainly based on feather color, raised by multiple farmers, either professional (MAG) or amateurs (MAR). We observed large differentiation between breeders (Fst=0.15) which was mainly due to separation between MAG and MAR (Fst=0.16). The differentiation between color varieties was stronger within the MAG breed (Fst=0.20) than in the MAR breed (0.12) despite having twice more varieties. This reflected a different way to raise their flocks with exchanges between breeders of the MAR population leading to more connectivity between varieties. This can be illustrated by the MNC color variety (Marans Noire Cuivrée) which is present into both professional and amateur farms but divided into two distinct groups according to the type of farm. Conversely, within the group of six amateur farms, individuals of the MNC variety clearly clustered together (fig. 7) whereas those of the MF variety (Marans Froment) did not. Thus the way of breeding, either professional or amateur, was the most prominent factor that shaped the genetic structure of the Marans breed. Exchanges between amateur breeders led to more genetic diversity conserved but varieties less differentiated from each other while the professional breeder led to more genetically distinguishable varieties. Some color varieties can be associated to variation in a few known pigmentation genes: *MC1R* for the red to black variation (Kerje et al., 2003), *CDKN2A* for sex-linked barring (Hellström et al., 2010) and *SLC45A2* for silver to gold variation (Gunnarsson et al., 2007). Yet, color varieties of the Marans breed appeared to be rather structured by management types. However all MAR varieties clustered together confirming that the breed concept remains meaningful even when different varieties are managed by a network of amateur breeders. This is of particular importance for future conservation plans of genetic resources since the scale to which monitor genetic diversity and the base unit to conserve are crucial for decision making.

### Which tools to evaluate management programs

#### On the interest of molecular tools for conservation

While Fis informed us about the management of populations given the allelic diversity present in the population, Fit allows for measuring the absolute genetic diversity present within the population as a deviation to the expectation with the approximate ancestral allele frequencies. Indeed we found a positive correlation between Fit (and a weaker one with Fis) and the proportion of fixed alleles similarly to what Muir et al. (2008) observed between the proportion of missing alleles and the inbreeding coefficient. This makes this Fit-like estimator a good index to figure out the genetic diversity conserved in populations. It is the result of both the effects of the size of the selection nucleus, the founder effect and the mating plans. In the present study, the mean value of Fit was 0.26 and it ranged from 0.11 to 0.46. These values are very similar to the inbreeding coefficients estimated from the ROH, F-ROH, that ranged from 0.13 to 0.42, with a very large correlation between the two estimates. An advantage of ROH inbreeding estimates used in this study (i.e. computed with Plink) is that it does not rely on any assumption about allelic frequencies, and thus it is insensitive to the number of individuals or populations considered contrary to Fit that needs allele frequencies in multiple populations. It makes ROH estimates an index of choice to monitor the genetic diversity of populations, in particular in absence of any additional information like pedigrees or accurate estimates of allelic frequencies which is often the case with small local populations. In addition, the length of ROH was also informative about the efficiency of the management program. Indeed for the local breeds experiencing such a program (i.e. group 1) we observed a significant and negative correlation between the mean length of ROH and the number of generations elapsed since the start of the program (figure 6), meaning less recent inbreeding for breeds that benefited from a management program early on, and thus efficient mating plans. Of course, the length of the ROH was also influenced by the number of founders at the start of the program. The trend observed with ROH was also observed with Ne estimates based on either pedigrees or linkage disequilibrium confirming their potential for the monitoring of genetic diversity of these populations as shown in previous studies (Doublet et al., 2019). A huge advantage of ROH is its simplicity and rapidity to compute, even without any information about allelic frequencies, which makes it possible to obtain individual estimates even without information on numerous individuals. In addition, Caballero et al. (2021) showed that ROH are very accurate for estimating inbreeding depression in populations with limited effective population size, which is often the case with local breeds. Nevertheless the weak and almost all non-significant correlations of Fis with other estimates, indicated that it brought original and unique information about the mating plan and inbreeding management, not provided by the other indices, either demographic or molecular.

The correlations found between molecular indicators and demographic parameters showed that the number of female founders was the most correlated with the mean length of ROH, slightly more than was the number of generations, and even more than was the number of male founders. This could be due to the fact that female founders are generally provided by many different farmers, each of them having a limited number of animals. Since the number of males is generally quite lower than the number of females, not all farmers provide a male but most farmers will provide a female, so that a high number of female founders indicates a larger genetic basis at the onset of the conservation program. Furthermore, a female chicken of a local breed can have from 50 to 100 progeny over one year if chicks are raised separately from the dams, showing a high potential for population expansion.

The fact that the number of generations in the program was strongly positively correlated with the genetic size estimated by LD (Ne_LD), and to a lesser extent with the genetic size estimated on pedigree data, is due to the fact that the most sustainable programs (higher number of generations) are those which better manage inbreeding. It occurs that these programs are also those which started with a higher number of females, as shown by the high correlation observed between Ne_LD and the number of founder females whereas the correlation between Ne_LD and the number of founder males was non significant. This means that the breeding program made possible to conserve most, if not all, female origins along the generations, which contributed more to maintain genetic diversity than did the number of founder males.

### Recommendations for conservation programs of small populations

Our results confirm that the size of the founder population is a critical parameter, but that a well planned management makes possible to maintain genetic diversity even in populations which started with a small number of founders. Considering that pedigree recording can be a daily constraint, we also show that molecular indicators can be used to efficiently monitor the conservation program at planned intervals checking for inbreeding and co-ancestry. For populations sub-divided, in many flocks for instance, López-Cortegano et al. (2019) suggested to also pay particular attention to haplotypic diversity. In addition, parentage assignation can also be conducted routinely for populations without control of mating (e.g. free range) to drive the selection of the reproducers for the next generation (i.e. RefGenDivA project). The spectacular growth of the number of reference genomes show that a subset of molecular markers could be available for a large number of animal species, either domestic or wild. It then becomes important to develop a standard and easy-to-use tool, proposed to all managers of small populations at a reasonable cost. This is the aim of the current development of two affordable multi-species SNP chip initiated in the H2020 IMAGE project (imageh2020.eu) for cattle, sheep, goat, horse, pig, chicken, buffalo, rabbit, quail, pigeon, duck and bee, with 10K markers for each species (Crooijmans et al., in prep.). Using the same genotyping tool will make it possible to compare genetic diversity and monitor conservation programs across populations and countries for a coordinated management of genetic resources at a larger scale than the country or the region. Muir *et al*. (2008) proposed to combine different chicken breeds to limit the loss of genetic diversity even if diversity recovery could be limited. They suggested that combining other breeds than commercial pure lines is a promising solution in terms of genetic conservation but we showed here that the breed structure and breed history, shaped by farmers’ practices, should be considered in order to identify the most relevant combinations and elements to conserve. Indeed Gicquel et al. (2020) showed that breeds benefiting from conservation programs aiming at increasing their population size did not always translate into an increase of their genetic variability. Thus explicit consideration of this genetic variability and identification of its determinants could help to make such conservation programs more efficient in the future.

## Conclusion

We characterized the high level of diversity in a large set of French local chicken breeds and thus their potential as valuable genetic resources for the future. Indeed such a large genetic diversity is an obligatory feature to cope with global changes and achieve a more sustainable production through a better adaptation to variable environmental conditions. We also showed that an appropriate management / breeding program can reconcile moderate production with conservation of genetic diversity both within and between breeds, while these are generally opposed. Indeed, avoiding genetic relatedness in the mating plan limited the increase of inbreeding, as frequently observed in other fancy breeds, while maintaining numerous breeds over the country. This was due to the particular French niche market, associated with a strong local appropriation of breeds to a territory and based on top quality products, often raised under free-range conditions for which selection pressure was weak. We revealed the importance of sampling a large number of female founders and the conservation of their subsequent families combined with a quick increase of the population size. This way one can ensure to conserve a maximum amount of the initial genetic diversity. Finally we stressed the usefulness and accuracy of ROH based estimates for evaluation and routine monitoring of genetic diversity, in particular in the absence of complete pedigrees or when only few samples are available which is often the case for local breeds.

## Acknowledgement

This project named BioDivA (Project 1258 - “BIODIVA Objectif PRM: caractérisation de la biodiversité des races locales de volailles françaises pour accompagner la mise en place du dispositif européen Protections des Races Menacées pour les Volailles”) was funded by CASDAR program of the French Agriculture Ministry. Financial support for genotyping and biobanking was provided by the CRB-Anim infrastructure project, ANR-11-INBS-0003, funded by the French National Research Agency in the frame of the ‘Investing for the Future’ program. Additional fundings for genotyping were obtained through the RefGenDivA project and the Rhones-Alpes region. We would like to thank the breed associations who permitted us to have access to the data. Finally we would like to acknowledge François Seigneurin, Maryse Boulay and Hervé Chapuis from the SYSAAF for providing and managing pedigree data.

## Data accessibility

Genomic data are available in the diversity browser of the IMAGE project (http://www.imageh2020.eu/) and will be also available on dryade.org (doi).

## Authors Contribution

MTB, DG and SL designed the project. MTB,AV, DG & FP organized data collection with the breeders and SYSAAF. GR designed the study, carried out the analysis and wrote the first draft of the manuscript. FP, DG, RR and SBF provided demographic data. GR, XR and MTB jointly supervised the study and contributed to the writing of the manuscript. All authors revised and approved the final version.

**Supplementary material S1.**
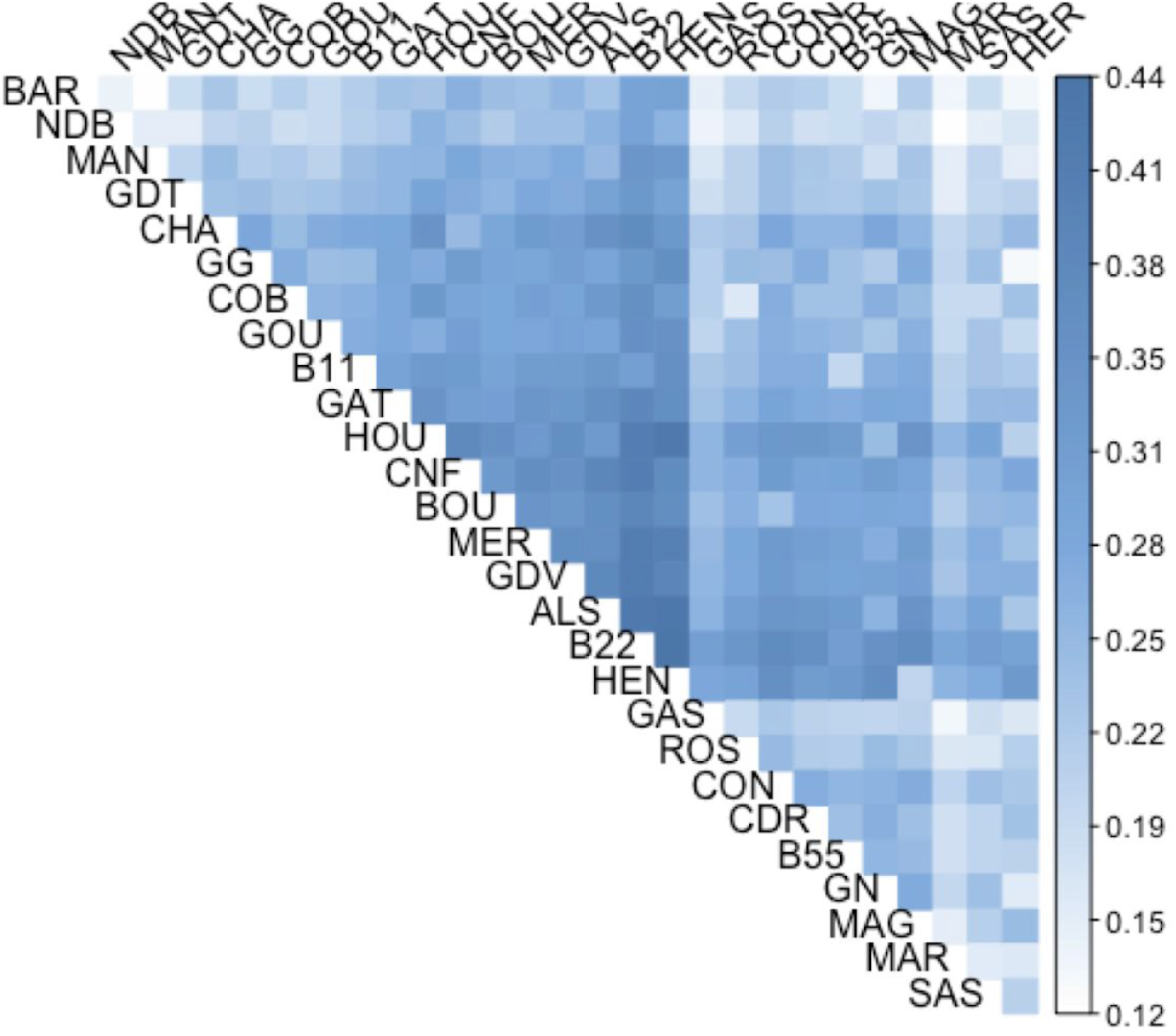
Matrix of pairwise Fst between populations.

**Supplementary material S2.**
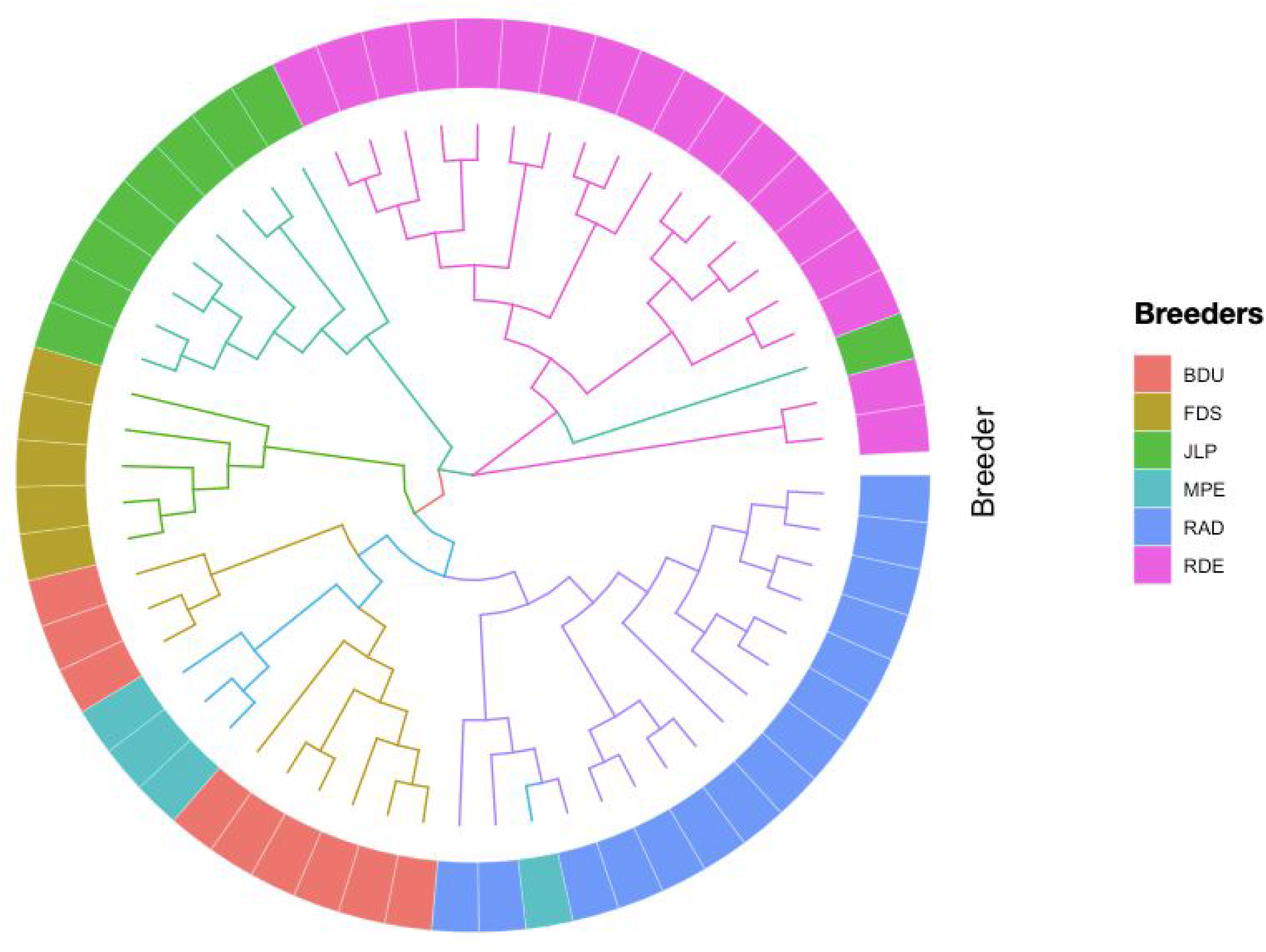
Unrooted neighbor-joining tree of the Hergnie breed. The colors of represent the different breeders.

**Supplementary material S3.**
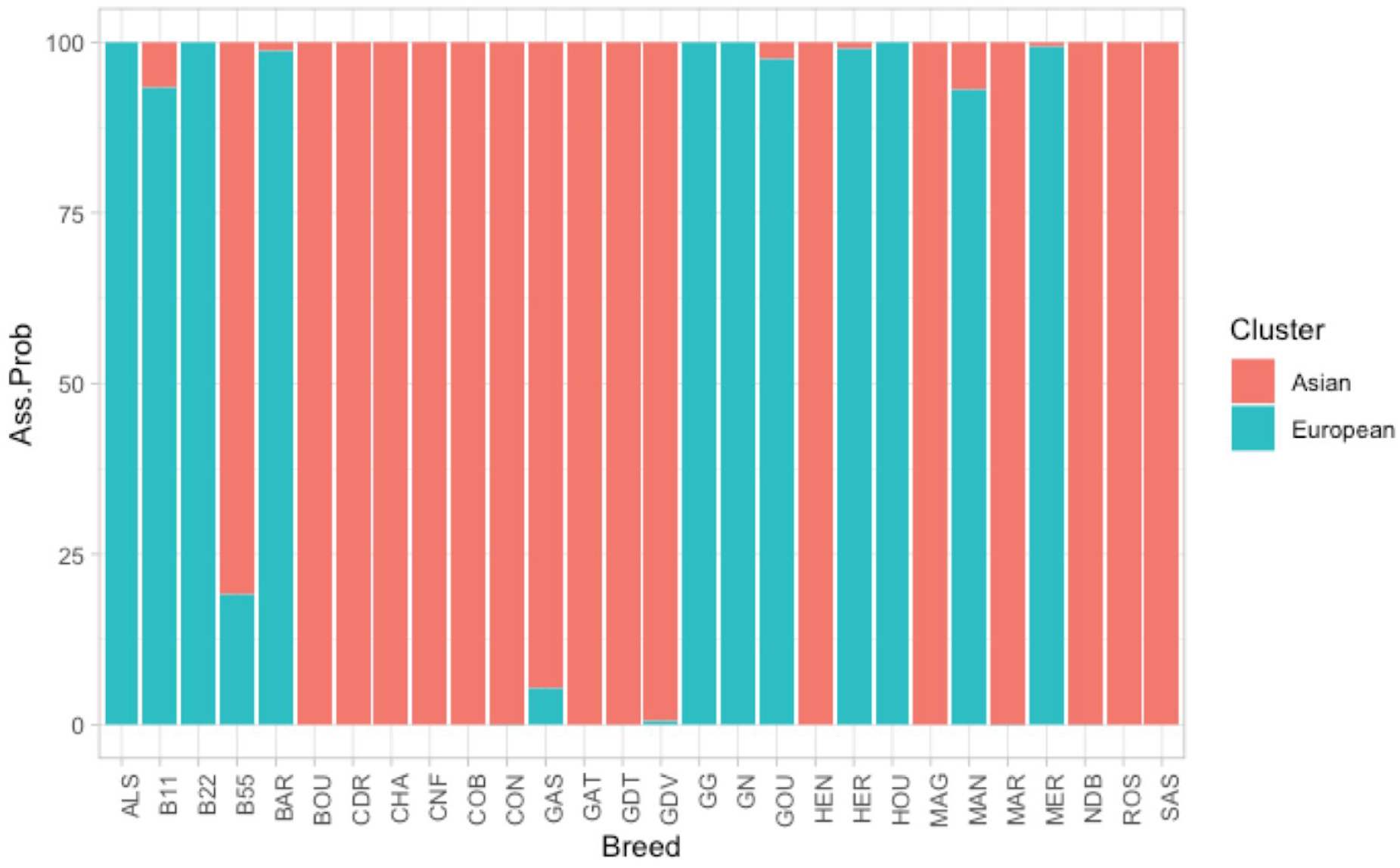
Mean assignation probability of populations to each of the 2 clusters, namely the Asian or European one.

**Supplementary material S4.** Within population genetic diversity molecular estimates.

| Breed | Fit | Fis | H_obs | Fixed (%) | F.ROH | Number.ROH | Length.ROH (bp) | Ne_LD |
| --- | --- | --- | --- | --- | --- | --- | --- | --- |
| NDB | 0.11 | 0.06 | 0.32 | 0.05 | 0.13 | 36.57 | 3180.87 | 75.8 |
| FG1 | 0.12 | 0 | 0.33 | 0.09 | 0.18 | 70.45 | 2308.59 | NA |
| FG2 | 0.14 | -0.02 | 0.35 | 0.09 | 0.16 | 60.6 | 2451.06 | NA |
| BAR | 0.15 | -0.02 | 0.35 | 0.09 | 0.13 | 42 | 2908.66 | 123.1 |
| GAS | 0.15 | -0.02 | 0.34 | 0.09 | 0.15 | 42.35 | 3198.61 | 93.8 |
| SGB | 0.15 | -0.02 | 0.34 | 0.1 | 0.15 | 58.01 | 2376.33 | NA |
| GDT | 0.19 | 0.02 | 0.32 | 0.13 | 0.18 | 66.83 | 2431.51 | 201.3 |
| MAR | 0.22 | 0.05 | 0.32 | 0.05 | 0.2 | 56.58 | 3138.85 | 33.1 |
| CDR | 0.23 | -0.01 | 0.34 | 0.17 | 0.21 | 77.95 | 2535.37 | 141 |
| B55 | 0.23 | -0.02 | 0.36 | 0.17 | 0.2 | 84.21 | 2209.78 | NA |
| CHA | 0.24 | -0.02 | 0.35 | 0.18 | 0.23 | 75.7 | 2789.59 | 41.5 |
| MAN | 0.24 | 0.08 | 0.31 | 0.15 | 0.23 | 43 | 4698.18 | 2.5 |
| GOU | 0.25 | -0.02 | 0.33 | 0.17 | 0.21 | 75.07 | 2652.41 | 173 |
| GN | 0.25 | -0.02 | 0.35 | 0.17 | 0.16 | 63.74 | 2353.95 | 86.2 |
| GG | 0.27 | -0.01 | 0.34 | 0.22 | 0.2 | 77.8 | 2425.79 | 1381.7 |
| B11 | 0.28 | -0.02 | 0.35 | 0.19 | 0.24 | 94.57 | 2363.01 | 1840.2 |
| CON | 0.29 | -0.01 | 0.34 | 0.19 | 0.28 | 62 | 4098.85 | 17.2 |
| HER | 0.3 | 0.08 | 0.32 | 0.07 | 0.23 | 50.05 | 4017.7 | 7 |
| GAT | 0.3 | -0.02 | 0.34 | 0.22 | 0.28 | 80.95 | 3236.04 | 171.4 |
| BOU | 0.32 | -0.03 | 0.35 | 0.22 | 0.28 | 100.23 | 2579.77 | 151.1 |
| CNF | 0.32 | -0.03 | 0.35 | 0.26 | 0.32 | 91.58 | 3235.62 | 48.7 |
| MAG | 0.33 | 0.05 | 0.31 | 0.16 | 0.31 | 104.31 | 2710.12 | 12.5 |
| MER | 0.34 | -0.02 | 0.35 | 0.29 | 0.31 | 84.95 | 3408.76 | 29.8 |
| LAY | 0.35 | -0.01 | 0.36 | 0.28 | 0.36 | 124.12 | 2697.89 | NA |
| **GDV** | 0.35 | -0.02 | 0.35 | 0.29 | 0.3 | 92.2 | 3024.89 | 122.7 |
| **HOU** | 0.38 | -0.01 | 0.35 | 0.3 | 0.24 | 89.71 | 2517.68 | 73.4 |
| **ALS** | 0.41 | -0.02 | 0.35 | 0.34 | 0.31 | 102.43 | 2803.78 | 26.3 |
| **B22** | 0.46 | -0.02 | 0.34 | 0.38 | 0.42 | 128.4 | 3029.55 | 644.3 |

## References

AR4 Climate Change 2007: Synthesis Report — IPCC, n.d. URL https://www.ipcc.ch/report/ar4/syr/ (accessed 1.7.21).

Berthouly, C., Bed’Hom, B., Tixier-Boichard, M., Chen, C.F., Lee, Y.P., Laloë, D., Legros, H., Verrier, E., Rognon, X., 2008. Using molecular markers and multivariate methods to study the genetic diversity of local European and Asian chicken breeds. Anim. Genet. 39, 121–129. https://doi.org/10.1111/j.1365-2052.2008.01703.x

Bodzsar, N., Eding, H., Revay, T., Hidas, A., Weigend, S., 2009. Genetic diversity of Hungarian indigenous chicken breeds based on microsatellite markers. Anim. Genet. 40, 516–523. https://doi.org/10.1111/j.1365-2052.2009.01876.x

Bortoluzzi, C., Crooijmans, R.P.M.A., Bosse, M., Hiemstra, S.J., Groenen, M.A.M., Megens, H.-J., 2018. The effects of recent changes in breeding preferences on maintaining traditional Dutch chicken genomic diversity. Heredity 121, 564–578. https://doi.org/10.1038/s41437-018-0072-3

Brandenburg, J.-T., Mary-Huard, T., Rigaill, G., Hearne, S.J., Corti, H., Joets, J., Vitte, C., Charcosset, A., Nicolas, S.D., Tenaillon, M.I., 2017. Independent introductions and admixtures have contributed to adaptation of European maize and its American counterparts. PLOS Genet. 13, e1006666. https://doi.org/10.1371/journal.pgen.1006666

Caballero, A., Villanueva, B., Druet, T., 2021. On the estimation of inbreeding depression using different measures of inbreeding from molecular markers. Evol. Appl. 14, 416–428. https://doi.org/10.1111/eva.13126

Cervantes, I., Goyache, F., Molina, A., Valera, M., Gutiérrez, J.P., 2011. Estimation of effective population size from the rate of coancestry in pedigreed populations. J. Anim. Breed. Genet. 128, 56–63. https://doi.org/10.1111/j.1439-0388.2010.00881.x

Chang, C.C., Chow, C.C., Tellier, L.C., Vattikuti, S., Purcell, S.M., Lee, J.J., 2015. Second-generation PLINK: rising to the challenge of larger and richer datasets. GigaScience 4. https://doi.org/10.1186/s13742-015-0047-8

Crooijmans, R.P.M.A., Gonzalez-Prendes, Rayner, Michele Tixier-Boichard, Michèle, in prep. Managing genetic diversity to ensure resilience using the IMAGE multi-species SNP arrays.

Dávila, S.G., Gil, M.G., Resino-Talaván, P., Campo, J.L., 2009. Evaluation of diversity between different Spanish chicken breeds, a tester line, and a White Leghorn population based on microsatellite markers. Poult. Sci. 88, 2518–2525. https://doi.org/10.3382/ps.2009-00347

Denis, B., 2017. Les races de poules. Formation, évolution, présentation générale. Rev. D’ethnoécologie.

Dixon, P., 2003. VEGAN, a package of R functions for community ecology. J. Veg. Sci. 14, 927–930. https://doi.org/10.1111/j.1654-1103.2003.tb02228.x

Do, C., Waples, R.S., Peel, D., Macbeth, G.M., Tillett, B.J., Ovenden, J.R., 2014. NeEstimator v2: re-implementation of software for the estimation of contemporary effective population size (Ne) from genetic data. Mol. Ecol. Resour. 14, 209–214. https://doi.org/10.1111/1755-0998.12157

Doublet, A.-C., Croiseau, P., Fritz, S., Michenet, A., Hozé, C., Danchin-Burge, C., Laloë, D., Restoux, G., 2019. The impact of genomic selection on genetic diversity and genetic gain in three French dairy cattle breeds. Genet. Sel. Evol. 51, 52. https://doi.org/10.1186/s12711-019-0495-1

Fairfull, R.W. (Agriculture C.R.S., Gowe, R.S., 1990. Genetics of egg production in chickens. Dev. Anim. Vet. Sci. Neth.

Fu, W., Dekkers, J.C., Lee, W.R., Abasht, B., 2015. Linkage disequilibrium in crossbred and pure line chickens. Genet. Sel. Evol. 47, 11. https://doi.org/10.1186/s12711-015-0098-4

Gerber, P.J., Steinfeld, H., Henderson, B., Mottet, A., Opio, C., Dijkman, J., Falcucci, A., Tempio, G., 2013. Tackling climate change through livestock: a global assessment of emissions and mitigation opportunities. Tackling Clim. Change Livest. Glob. Assess. Emiss. Mitig. Oppor.

Gicquel, E., Boettcher, P., Besbes, B., Furre, S., Fernández, J., Danchin-Burge, C., Berger, B., Baumung, R., Feijóo, J.R.J., Leroy, G., 2020. Impact of conservation measures on demography and genetic variability of livestock breeds*. Animal 14, 670–680. https://doi.org/10.1017/S1751731119002672

Granevitze, Z., Hillel, J., Chen, G.H., Cuc, N.T.K., Feldman, M., Eding, H., Weigend, S., 2007. Genetic diversity within chicken populations from different continents and management histories. Anim. Genet. 38, 576–583. https://doi.org/10.1111/j.1365-2052.2007.01650.x

Groenen, M.A., Megens, H.-J., Zare, Y., Warren, W.C., Hillier, L.W., Crooijmans, R.P., Vereijken, A., Okimoto, R., Muir, W.M., Cheng, H.H., 2011. The development and characterization of a 60K SNP chip for chicken. BMC Genomics 12, 274. https://doi.org/10.1186/1471-2164-12-274

Gunnarsson, U., Hellström, A.R., Tixier-Boichard, M., Minvielle, F., Bed’hom, B., Ito, S., Jensen, P., Rattink, A., Vereijken, A., Andersson, L., 2007. Mutations in SLC45A2 Cause Plumage Color Variation in Chicken and Japanese Quail. Genetics 175, 867–877. https://doi.org/10.1534/genetics.106.063107

Harper, G.C., Makatouni, A., 2002. Consumer perception of organic food production and farm animal welfare. Br. Food J. 104, 287–299. https://doi.org/10.1108/00070700210425723

Hellström, A.R., Sundström, E., Gunnarsson, U., Bed’Hom, B., Tixier-Boichard, M., Honaker, C.F., Sahlqvist, A.-S., Jensen, P., Kämpe, O., Siegel, P.B., Kerje, S., Andersson, L., 2010. Sex-linked barring in chickens is controlled by the CDKN2A /B tumour suppressor locus. Pigment Cell Melanoma Res. 23, 521–530. https://doi.org/10.1111/j.1755-148X.2010.00700.x

Hoffman, A.J., 2010. Climate Change as a Cultural and Behavioral Issue: Addressing Barriers and Implementing Solutions (SSRN Scholarly Paper No. ID 2933572). Social Science Research Network, Rochester, NY. https://doi.org/10.2139/ssrn.2933572

Howden, S.M., Soussana, J.-F., Tubiello, F.N., Chhetri, N., Dunlop, M., Meinke, H., 2007. Adapting agriculture to climate change. Proc. Natl. Acad. Sci. 104, 19691–19696. https://doi.org/10.1073/pnas.0701890104

Huson, D.H., Bryant, D., 2006. Application of phylogenetic networks in evolutionary studies. Mol. Biol. Evol. 23, 254–267. https://doi.org/10.1093/molbev/msj030

Jombart, T., 2008. adegenet: a R package for the multivariate analysis of genetic markers. Bioinformatics 24, 1403–1405. https://doi.org/10.1093/bioinformatics/btn129

Jombart, T., Devillard, S., Balloux, F., 2010. Discriminant analysis of principal components: a new method for the analysis of genetically structured populations. BMC Genet. 11, 94. https://doi.org/10.1186/1471-2156-11-94

Kerje, S., Lind, J., Schütz, K., Jensen, P., Andersson, L., 2003. Melanocortin 1-receptor (MC1R) mutations are associated with plumage colour in chicken. Anim. Genet. 34, 241–248. https://doi.org/10.1046/j.1365-2052.2003.00991.x

López-Cortegano, E., Pouso, R., Labrador, A., Pérez-Figueroa, A., Fernández, J., Caballero, A., 2019. Optimal Management of Genetic Diversity in Subdivided Populations. Front. Genet. 10. https://doi.org/10.3389/fgene.2019.00843

Malomane, D.K., Simianer, H., Weigend, A., Reimer, C., Schmitt, A.O., Weigend, S., 2019. The SYNBREED chicken diversity panel: a global resource to assess chicken diversity at high genomic resolution. BMC Genomics 20, 345. https://doi.org/10.1186/s12864-019-5727-9

McQuillan, R., Leutenegger, A.-L., Abdel-Rahman, R., Franklin, C.S., Pericic, M., Barac-Lauc, L., Smolej-Narancic, N., Janicijevic, B., Polasek, O., Tenesa, A., MacLeod, A.K., Farrington, S.M., Rudan, P., Hayward, C., Vitart, V., Rudan, I., Wild, S.H., Dunlop, M.G., Wright, A.F., Campbell, H., Wilson, J.F., 2008. Runs of Homozygosity in European Populations. Am. J. Hum. Genet. 83, 359–372. https://doi.org/10.1016/j.ajhg.2008.08.007

Muir, W.M., Wong, G.K.-S., Zhang, Y., Wang, J., Groenen, M.A.M., Crooijmans, R.P.M.A., Megens, H.-J., Zhang, H., Okimoto, R., Vereijken, A., Jungerius, A., Albers, G.A.A., Lawley, C.T., Delany, M.E., MacEachern, S., Cheng, H.H., 2008. Genome-wide assessment of worldwide chicken SNP genetic diversity indicates significant absence of rare alleles in commercial breeds. Proc. Natl. Acad. Sci. 105, 17312–17317. https://doi.org/10.1073/pnas.0806569105

Notter, D.R., 1999. The importance of genetic diversity in livestock populations of the future1. J. Anim. Sci. 77, 61–69. https://doi.org/10.2527/1999.77161x

Paradis, E., Schliep, K., 2019. ape 5.0: an environment for modern phylogenetics and evolutionary analyses in R. Bioinformatics 35, 526–528. https://doi.org/10.1093/bioinformatics/bty633

Parry, M.L., 2019. Climate Change and World Agriculture. Routledge.

Pembleton, L.W., Cogan, N.O.I., Forster, J.W., 2013. StAMPP: an R package for calculation of genetic differentiation and structure of mixed-ploidy level populations. Mol. Ecol. Resour. 13, 946–952. https://doi.org/10.1111/1755-0998.12129

Purcell, S., Neale, B., Todd-Brown, K., Thomas, L., Ferreira, M.A.R., Bender, D., Maller, J., Sklar, P., de Bakker, P.I.W., Daly, M.J., Sham, P.C., 2007. PLINK: A Tool Set for Whole-Genome Association and Population-Based Linkage Analyses. Am. J. Hum. Genet. 81, 559–575.

Qu, L., Li, X., Xu, Guifang, Chen, K., Yang, H., Zhang, L., Wu, G., Hou, Z., Xu, Guiyun, Yang, N., 2006. Evaluation of genetic diversity in Chinese indigenous chicken breeds using microsatellite markers. Sci. China C Life Sci. 49, 332–341. https://doi.org/10.1007/s11427-006-2001-6

Tixier-Boichard, M., Bed’hom, B., Rognon, X., 2011. Chicken domestication: From archeology to genomics. C. R. Biol., On the trail of domestications, migrations and invasions in agriculture 334, 197–204. https://doi.org/10.1016/j.crvi.2010.12.012

Verrier, E., Tixier-Boichard, M., Bernigaud, R., Naves, M., 2005. Conservation and value of local livestock breeds: usefulness of niche products and/or adaptation to specific environments. Anim. Genet. Resour. Inf. 36, 21–31. https://doi.org/10.1017/S1014233900005538

Wei, T., Simko, V., Levy, M., Xie, Y., Jin, Y., Zemla, J., 2017. corrplot: Visualization of a Correlation Matrix.

Wickham, H., 2011. ggplot2. WIREs Comput. Stat. 3, 180–185. https://doi.org/10.1002/wics.147

Wilkinson, S., Wiener, P., Teverson, D., Haley, C.S., Hocking, P.M., 2012. Characterization of the genetic diversity, structure and admixture of British chicken breeds. Anim. Genet. 43, 552–563. https://doi.org/10.1111/j.1365-2052.2011.02296.x

Windhorst, H.-W., 2006. Changes in poultry production and trade worldwide. Worlds Poult. Sci. J. 62, 585–602. https://doi.org/10.1017/S0043933906001140

Yu, G., Smith, D.K., Zhu, H., Guan, Y., Lam, T.T.-Y., 2017. ggtree: an r package for visualization and annotation of phylogenetic trees with their covariates and other associated data. Methods Ecol. Evol. 8, 28–36. https://doi.org/10.1111/2041-210X.12628

Zhang, J., Nie, C., Li, X., Ning, Z., Chen, Y., Jia, Y., Han, J., Wang, L., Lv, X., Yang, W., Qu, L., 2020. Genome-Wide Population Genetic Analysis of Commercial, Indigenous, Game, and Wild Chickens Using 600K SNP Microarray Data. Front. Genet. 11. https://doi.org/10.3389/fgene.2020.543294

